# Liver Organoid and T Cell Coculture Models Cytotoxic T Cell Responses against Hepatitis C Virus

**DOI:** 10.1101/2021.08.10.455738

**Authors:** Vaishaali Natarajan, Camille R Simoneau, Ann L Erickson, Nathan L Meyers, Jody L Baron, Stewart Cooper, Todd C McDevitt, Melanie Ott

## Abstract

Hepatitis C virus (HCV) remains a global public health challenge with an estimated 71 million people chronically infected, with surges in new cases and no effective vaccine. New methods are needed to study the human immune response to HCV since *in vivo* animal models are limited and *in vitro* cancer cell models often show dysregulated immune and proliferative responses. Here we developed a CD8^+^ T cell and adult stem cell liver organoid system using a microfluidic chip to coculture 3D human liver organoids embedded in extracellular matrix with HLA-matched primary human T cells in suspension. We then employed automated phase contrast and immunofluorescence imaging to monitor T cell invasion and morphological changes in the liver organoids. This microfluidic coculture system supports targeted killing of liver organoids when pulsed with a peptide specific for HCV nonstructural protein 3 (NS3) (KLVALGINAV) in the presence of patient-derived CD8^+^ T cells specific for KLVALGINAV. This demonstrates the novel potential of the coculture system to molecularly study adaptive immune responses to HCV in an *in vitro* setting using primary human cells.

## Introduction

Hepatitis C virus (HCV) is a positive-sense single-stranded RNA virus which targets hepatocytes, usually establishes chronic infection and, untreated, can progress to cirrhosis and hepatocellular carcinoma^1^. About 20% of patients clear infections spontaneously, but the remainder establish a lifelong infection, which is often not diagnosed until severe liver disease has occurred^2^. While the incidence of HCV in the United States has remained lower than in some other regions, the opioid epidemic has led to an increase in HCV cases due to needle sharing^3^. Despite the recent introduction of direct-acting antivirals against HCV, the global disease burden remains high, related to lack of access to treatment, expense of drugs, and the possibility of reinfection after successful treatment is completed^4^. In addition, despite a sustained virologic response (SVR), the risk of hepatocellular carcinoma in patients treated when advanced fibrosis was present remains higher in previously HCV-infected individuals compared to those who have never been infected^5^. This underscores the need for an effective vaccine, the development of which has been hampered by the high mutation rate of the virus and the lack of broadly neutralizing antibody induction after natural infection or vaccination strategies^6^.

HCV demonstrates limited tropism and can productively infect chimpanzees and humans only^7^. The dynamics of the immune response to HCV have been well studied in chimpanzee models, which show that CD8+ and CD4+ T cells play an important role in the control of HCV infection^2,8^. Although anti-HCV antibodies are produced, they do not protect against reinfection, and notable is that patients with congenital agammaglobulinemia have been able to spontaneously resolve acute hepatitis C^9^. However, the most recent human vaccine trial that succeeded in eliciting anti-HCV T cell responses did not offer any protection against chronic HCV infection^10^.

With chimpanzees no longer available for animal studies, there is an urgent need for laboratory models of HCV immunity. However, physiologically relevant laboratory models of HCV have been challenging to establish as primary human hepatocytes do not sustain long-term culture due to rapid dedifferentiation *in vitro*, and thus cannot be chronically infected with HCV^11^. Therefore, most research has relied on viral strains that can robustly replicate in various subclones of the Huh7 hepatoma cell lines^12-14^. However, the genetic perturbations in the antiviral interferon response that make Huh-7 cells highly permissive to HCV infection, also result in dysfunctional innate immune defenses, preventing accurate modeling of HCV immunity^12^. In addition, Huh7 cells lack proper cellular polarity and therefore demonstrate a non-physiological distribution of the surface receptors exploited by HCV for cell entry^12^.

Exciting recent developments in organoid technologies have enabled the creation of liver organoids from pluripotent and adult stem cells that replicate key structural and functional features of the organ and therefore can be employed for modeling HCV infection in the liver. Adult stem cells (ASCs) derived from the liver can be grown in a defined culture condition to robustly generate 3D liver organoids that can be maintained in culture without developing a senescent phenotype for multiple months^15-17^. Long-term culture of liver organoids can be initiated through selection of epithelial cellular adhesion molecule (EpCAM)^+^ cells from liver biopsies because the EpCAM+ compartment of the liver cells is enriched for ASCs.

ASC-derived liver organoids retain cell polarity rendering them highly suitable for the study of HCV entry and replication^18^. Embryonic stem cell (ESC)- and induced pluripotent stem cell (iPSC)-derived hepatic organoids have been successfully infected with clones of JFH1, a genotype 2a HCV strain, and with primary isolates of HCV^19-21^. Since paired liver and blood samples can be obtained from ASC donors, ASC-derived organoids are well suited for the studies of host-pathogen interactions and the immune response to infection and cancer in an autologous coculture setting^22,23^.

Here, we employ a static microfluidic chip to emulate the physiological interaction between a solid tissue, the liver, and the T cells in circulation. ASC-derived liver organoids are embedded in extracellular matrix within the central channel of a microfluidic chip and the CD8^+^ T cells suspended in media in adjacent, inter-connected microfluidic channels. The fluidics configuration can be manipulated to drive the T cells from media towards the central channel where the liver organoids are contained. We generated an HCV A0201-KLVALGINAV (NS3 aa1406-1415) CD8^+^ T cell clone and matched it with a liver organoid donor to allow coculture. Coculture in the microfluidics chip enabled the real-time imaging to monitor the organoid/T cell interactions, and precise control over cellular interactions through modulation of culture parameters, including flow rate and soluble factor gradients. This coculture system, which recapitulates immune cell dynamics in liver tissue microenvironments, represents a powerful new tool to model critical cellular features of HCV immunity.

## Results

### 1. Adult stem cell-derived liver organoids express HLA class I

ASC-derived organoids were grown from liver samples obtained from clinical resection procedures, as previously described^15^. The resulting organoids consist mostly of a single epithelial cell layer surrounding a hollow center; in the stem-cell state, liver organoids can be maintained and expanded in basement membrane extract (BME) for at least four months without loss of viability^18^. Antigen presentation on the surface of a target cell is required for T cell interaction to initiate a primary immune response. Thus, to test whether liver organoids could function as antigen-presenting cells, the levels of the class I HLA markers, HLA-A, -B, -C, were measured by quantitative RT-PCR (Fig. 1A). We compared HLA expression in organoids obtained from HCV- and HCV+ donors, and observed no significant difference between the levels of expression based on viral status, as has been previously shown^24^ To determine the polarity of HLA class I expression, we used light sheet microscopy to examine the entire organoid (∼200 μm diameter) and stained for beta-2 microglobulin, a component of conformationally and functionally mature HLA class I complexes. These analyses indicated that beta-2 microglobulin was readily expressed and localized on the “outside”, basolateral surface of the organoids (Fig. 1B).

**Figure 1.**
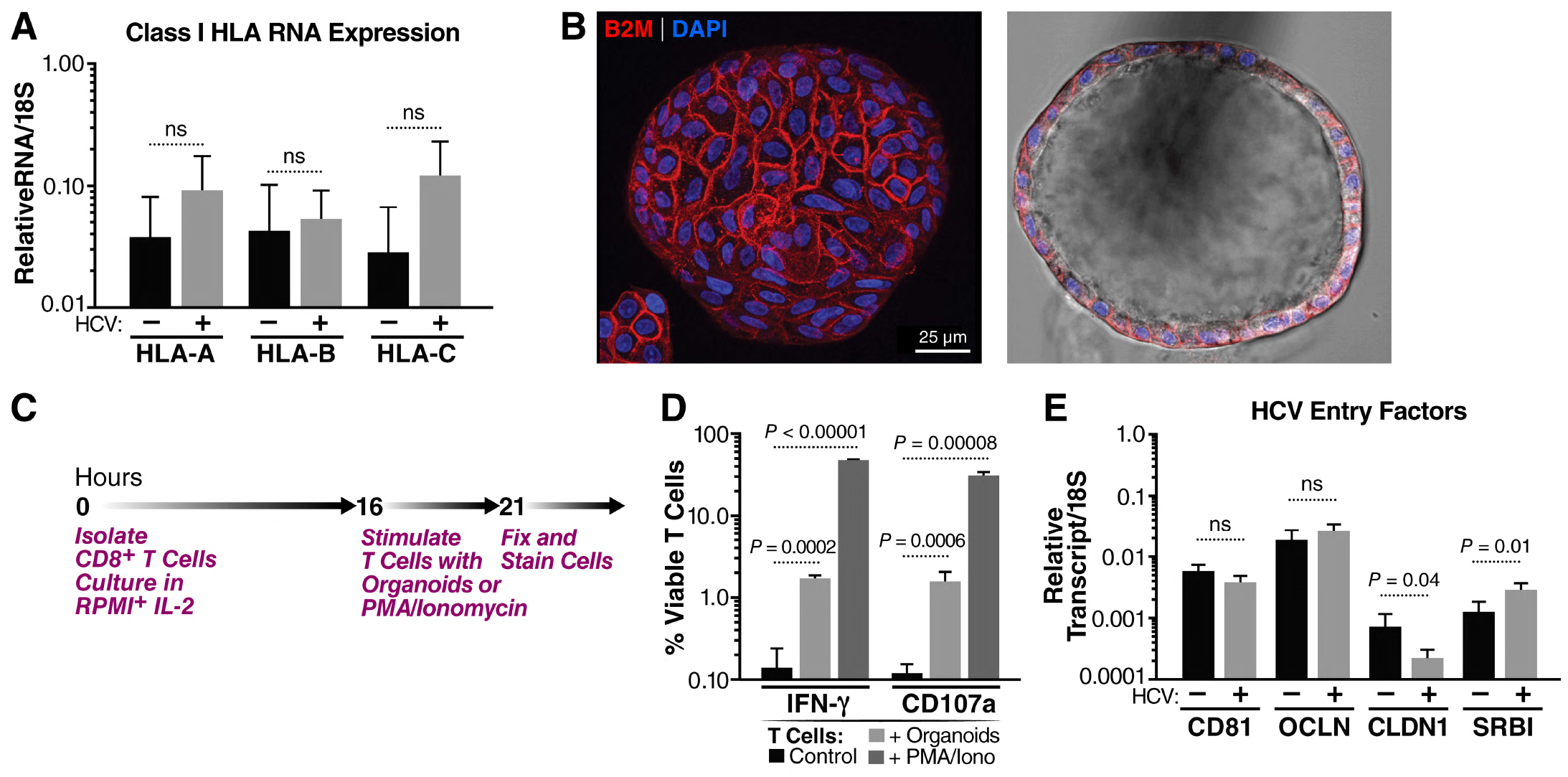
Human liver organoids in the stem cell (EM) state express the necessary factors to interact with CD8^+^ T cells and HCV virions. **(A)** Expression levels of HLA Class I genes as determined by RT-qPCR from HCV- (N=5) and HCV+ (N=4) liver organoids. RNA levels were standardized to the 18S gene and significance was calculated with an unpaired t-test. NS = not significant. **(B)** Representative light sheet images of a liver organoid stained with the pan-class I HLA marker β_2_ microglobulin (B2M; red) and DAPI (blue). **(C)** Schematic timeline showing the experimental protocol for T cell stimulation. **(D)** Expression of IFN-γ and CD107a on CD8^+^ T cells stimulated by PMA and ionomyacin or HLA-mismatched organoids as measured by flow cytometry and gating on viable CD8^+^ T cells. **(E)** Expression levels of HCV entry factors CD81, occludin (OCLN), claudin1 (CLDN1) and SRB1 by RT-qPCR from HCV- (N=4) and HCV+ (N=5) liver donors. All genes were standardized to 18S and significance was calculated with an unpaired t-test. NS = not significant.

Next, we tested whether CD8^+^ T cells could recognize and react to the HLA class I molecules on the surface of the organoids. CD8^+^ T cells were isolated from heathy blood donors and cultured overnight in the presence of interleukin-2 (IL-2) before incubating the T cells with allogenic organoids for 5 hours in a well-based assay without embedding the organoids in BME (Fig. 1C). Incubation with the allogenic organoids induced robust expression of interferon-gamma (IFN-γ), a proinflammatory cytokine, and of CD107a, a degranulation marker (Fig. 1D). Induction by organoids was similar to the induction by phorbol esters (PMA) combined with ionomycin, which is a strong T cell-activating stimulus. Collectively, these observations indicate that the organoids act as efficient antigen-presenting cells for CD8^+^ T cells.

As HCV entry occurs via multiple surface-expressed receptors, we confirmed the expression of HCV receptors CD81, claudin, occludin and scavenger receptor B1 (SRB1) in liver organoids. No significant difference in CD81 or occludin transcript levels was observed between donors; however, claudin mRNA expression was on average higher and SRB1 lower in HCV-negative samples. (Fig 1E). Due to the expression of functional HLA class I and HCV receptors on the organoid cell surface, we conclude that these organoids have the required proteins to model T cell interactions in the context of an infection.

### 2. Generation and characterization of the HCV A0201-KLVALGINAV (NS3 aa1406-1415) CD8^+^ T cell clone

To model antigen-mediated killing of virally infected cells, we generated T cell clones that could specifically recognize distinct HCV peptide antigens presented by liver organoid cells (Fig 2A). To that end, the HCV NS3 protein contains numerous immunologically relevant epitopes, including the HLA-A0201-restricted peptide, KLVALGINAV (aa 1406–1415)^25,26^. We therefore isolated CD8^+^ T cells specific for the KLVALGINAV peptide from an individual who spontaneously resolved HCV. The antigen-specific T cell population was sorted from peripheral blood using an HLA-A0201-KLVALGINAV tetramer (Figure 2B). Individual CD8^+^ T cells clones were expanded in culture with irradiated allogeneic peripheral blood mononuclear cells (PBMC), anti-CD3 antibodies and recombinant IL-2. The expanded CD8^+^ T cell clones were assessed for antigen specificity in a chromium release assay against HLA-A0201-expressing targets pulsed with either the A0201-KLVALGINAV peptide or an HLA-A0201-restricted control peptide. A representative chromium release assay using one of the clones, SR01-78, is shown in Figure 2C (orange and grey lines). To confirm that clone SR01-78 also reacted to the endogenously processed KLVALGINAV antigen, the HLA-A0201-expressing target cells were infected with a recombinant vaccinia virus expressing the HCV NS3 (vvNS3) protein or with wild-type vaccinia virus (vvWT). This experiment demonstrated that clone SR01-78 reacted not only to exogenously loaded KLVALGINAV but also to the endogenously processed antigen (Fig. 2C, blue and purple lines).

**Figure 2.**
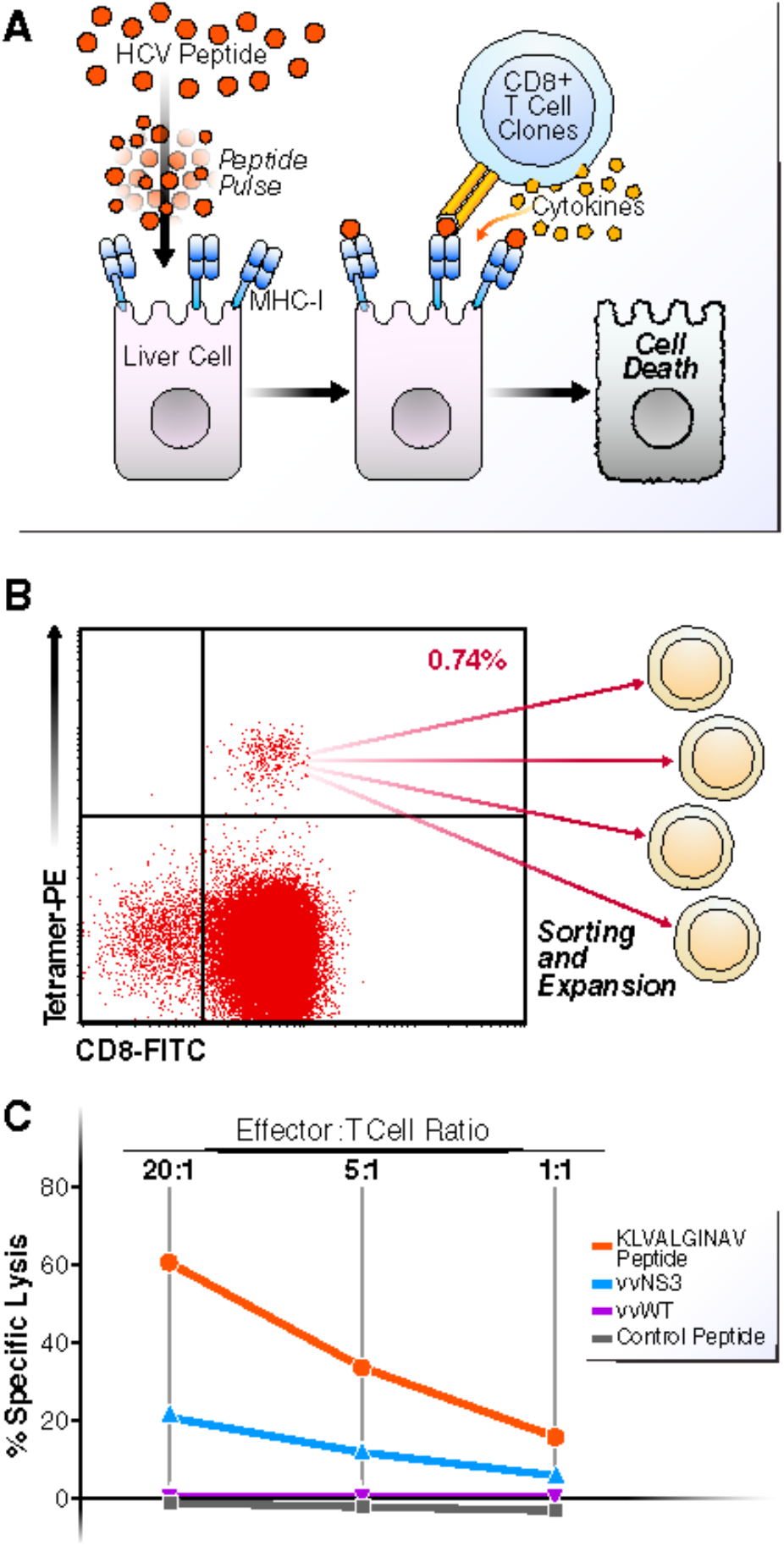
HCV antigen-specific lysis of A0201 B cell targets by KLVALGINAV CD8^+^ T cell clone SR01-78 from an individual with spontaneous resolved HCV. **(A)** Schematic detailing exogenous peptide pulsing of liver cells and stimulation of T cells. **(B)** Sorting strategy for HCV tetramer positive CD8^+^ T cells. **(C)** Chromium release assay demonstrating antigen specific lysis of A0201 class I MHC transfected B cell (721.221) exogenously loaded peptide targets (KLVALGINAV and control peptide) and endogenously synthesized antigen targets (vvNS3 and control vvWT) at effector to target ratios of 20:1, 5:1 and 1:1.

### 3. Culture of liver organoids and CD8^+^ T cells in a microfluidic chip

Previous reports have used simple well-based or droplet-based coculture of T cells with organoid cultures in a cancer immunity context as we have done in our pilot results in Figure 1^22^. Lack of BME embedment in the traditional well-based assay renders it unsuitable for long-term cultures of liver organoids as the extracellular matrix is essential for the maintenance of organoid structure, viability and polarity. On the other hand, the conventional droplet based coculture supports long-term organoid culture, but is unsuitable for the temporal study of individual organoids in response to HCV or the presence of immune cells. Therefore, to establish a tractable coculture of T cells and liver organoids that allows live monitoring of the interaction and potential killing over time, we used a commercially available microfluidic chip device (Fig. 3A)^27^. This contains a central channel where organoids can be embedded in basement membrane extract (BME), and flanking media channels on the top and bottom where media and T cells can be introduced. The content of the central channel remains contained while interstitial flow is permitted between the flanking media channels. A pressure difference is created and maintained between the media channels via differential media volume reservoirs connected to the channels to ensure transverse interstitial flow of media across the central channel. T cells added into the higher-pressure channel can thus be moved through the central channel by this transverse flow. This system therefore allows controlled entry of T cells into the central channel. Since the microfluidic chip also functions as a microscopy slide, it enables convenient tracking of T cell:organoid interactions via live, brightfield and fluorescence microscopy. We used phase contrast microscopy for tracking and morphological characterization of organoids, and fluorescence microscopy to quantify T cells labeled with a fluorescent dye (CellTracker Green).

**Figure 3.**
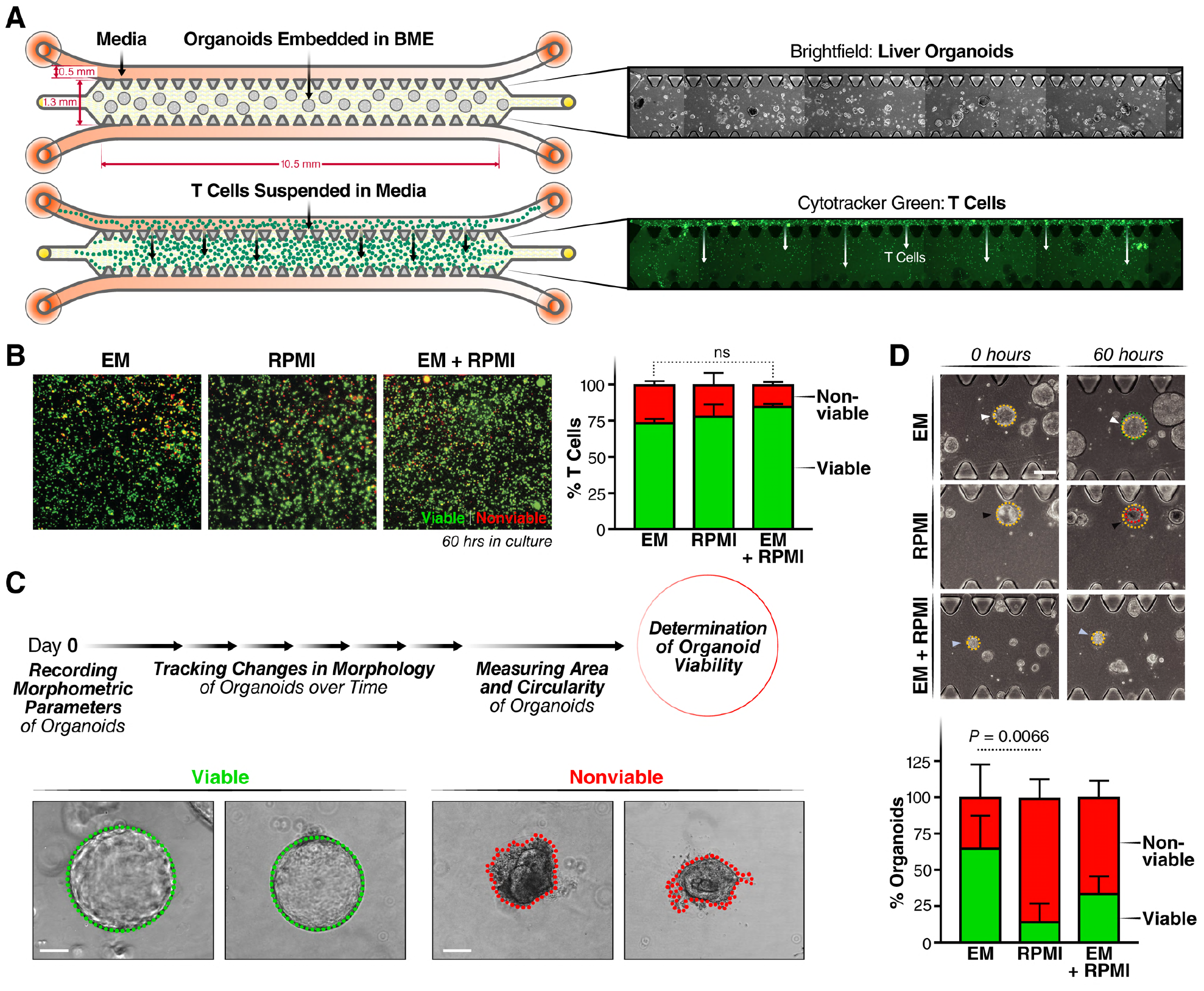
Establishing ideal culture conditions for T cells and Liver organoids in microfluidic chips. **(A)** Schematic detailing the experimental setup for monoculture of organoids and T cells, individually, in 3D microfluidic chips with corresponding microscopy images. The chip is made up of a central channel and 2 media channels on the top and bottom. Organoids are cultured in the middle channel in BME (see brightfield images, right). T cells are loaded into the chip from the media channels and migrate into the BME in the center of the chip driven by pressure gradient (see fluorescent images, right). T cells are tracked by CellTracker Green. **(B)** Quantification of T cell viability in RPMI, EM or RPMI:EM (1:1 volumetric ratio) media. T cells stained with CellTracker (green) are counted as viable cells and DRAQ7+ cells (red) are counted nonviable. **(C)** Organoids are classified as live or dead based on their intact morphology. Green outline (left) indicates healthy organoid with epithelial integrity and red outline (right) indicates atrophy and loss in viability. (**D**) Organoid survival in 3 media conditions: complete RPMI, EM and a 1:1 ratio of EM:complete RPMI. Brightfield images of the organoids at 12 and 60 hours in all media (top). Representative image of the classification of the viability of the organoids in each condition (left). Quantification was done on three replicates by measuring the area of the organoids. At 0 hours, the area is noted in yellow, and at the bottom the new area is noted with a circle in green (growth), yellow (same size) and red (smaller).

### 4. Optimization of coculture media conditions

T cells are typically cultured in RPMI mammalian cell culture medium supplemented with 10% fetal bovine serum (FBS). In contrast, liver organoids are cultured in Dulbecco’s Modified Eagle Medium (DMEM), which lacks several components of RPMI but also requires agonists for the Wnt and Notch signaling pathways to support organoid growth^15^. Importantly, organoids are usually cultured without the addition of FBS since the abundant soluble factor signals in FBS can trigger heterogenous differentiation of the ASC-derived liver organoids. Therefore, to determine which medium would be optimal for the coculture system, we tested the individual viabilities of T cells and liver organoids when monocultured in organoid medium (or EM), the T cell RPMI medium and a 1:1 mixture of both. We set up monocultures of either T cells or liver organoids in the microfluidic chips and monitored the viability of cells through microscopy. The viability of T cells was tracked by fluorescent imaging in the microfluidic device with the addition of DRAQ7 viability dye, which enables *in situ* labeling of nonviable cells. We quantified CellTracker^+^ or DRAQ7^+^ cells to calculate the fraction of viable or nonviable T cells, respectively. T cells maintained similar viabilities (>75%) in all three culture conditions over a 60-hour period with no statistically significant difference between the RPMI or EM media (Fig. 3B). For organoids, phase contrast imaging and morphological characterization were employed to quantify viability. Organoids with a spherical shape and good epithelial integrity were classified as viable, whereas liver organoids with irregular shape and loss of growth, as indicated by the red outline, were deemed nonviable organoids (Fig. 3C). After 60 hours in RPMI media, organoid viability dropped to below 25%, while remaining above 75% in the EM media. A 1:1 mix of both media remained below 50% viability. Thus, EM media was chosen for use in subsequent coculture studies as it supported both T cell and organoid survival.

### 5. Coculture of organoids and T cells

Once the optimal media conditions were determined in independent monocultures, the organoid:T cell cultures was cocultured in the microfluidic chip (Fig. 4A). Liver organoids under 100 µm in diameter were mixed with BME and added to the fenestrated central channel resulting in a spatially stable distribution of the organoids. The A0201-KLVALGINAV CD8^+^ T cells, pre-stained with Celltracker Green, were added to one of the flanking media channels in EM culture media followed by continuous media flow through both channels (Fig, 4A). T cells migrated through the gaps in the fenestrated central lane in the presence of a non-targeting peptide with numbers reproducibly increasing over the 60-hour observation period (Fig. 4B). Organoid viability in the central channel remained stable with a modest, but non-significant decrease over the 60 hour-time period. This indicates that T cell coculture and peptide pulsing are not inherently toxic to the liver organoids.

**Figure 4.**
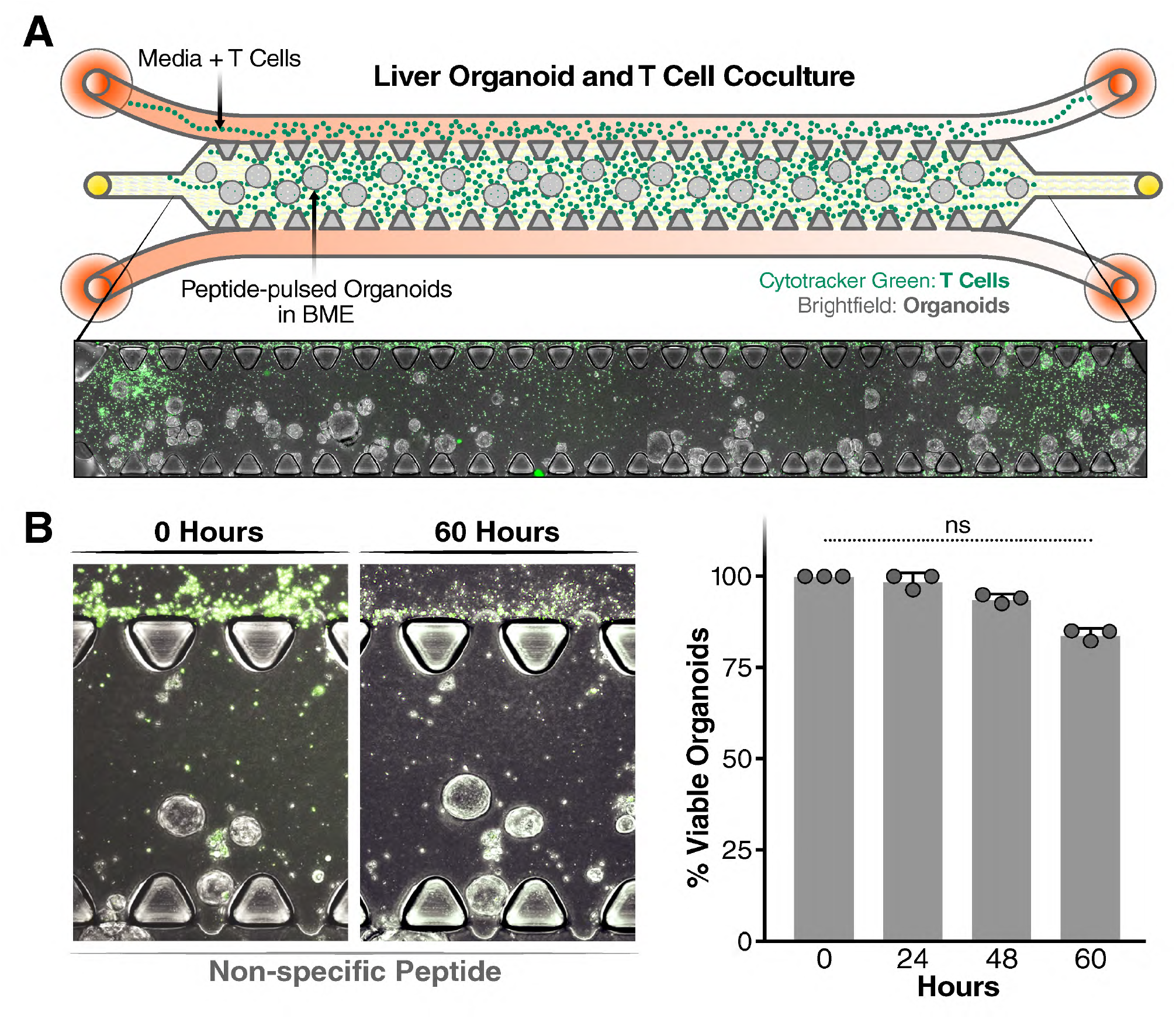
Coculture of T cells and liver organoids. **(A)** Schematic representation of the liver organoid and T cell coculture system (top) and representative image of the coculture system in the microfluidic chip (bottom). **(B)** Quantification of liver organoid viability after pulsing with peptide with representative images of the coculture (left) and quantification over 60 hours (right).

### 6. T cell response to HCV peptide-pulsed organoids

To determine whether the cloned T cells recognize and kill organoids expressing their cognate peptide, organoids were pulsed with the KLVALGINAV HCV-specific peptide and then loaded into the central channel, while A0201-KLVALGINAV-specific T cells were added in the media channel. Microscopy images were acquired at 15, 40 and 60 hours after the start of the coculture with representative images shown (Fig 5A). At 40 hours, peptide-pulsed organoids showed a more than 10-fold decrease in viability compared to the organoids not pulsed with the HCV peptide (p < 0.001) (Fig 5B). At 60 hours, 80% of the organoids were deemed non-viable in the peptide-pulsed conditions while only 15% cell death was recorded in the control condition (p < 0.001) (Fig. 5B). When we counted the number of T cells that had migrated into the central channel to confirm that the increase in cell death in the peptide-pulsed condition was not simply due to more T cells entering the central channel, we found a higher number of T cells in the control condition, excluding this possibility (Fig, 5C).

**Figure 5.**
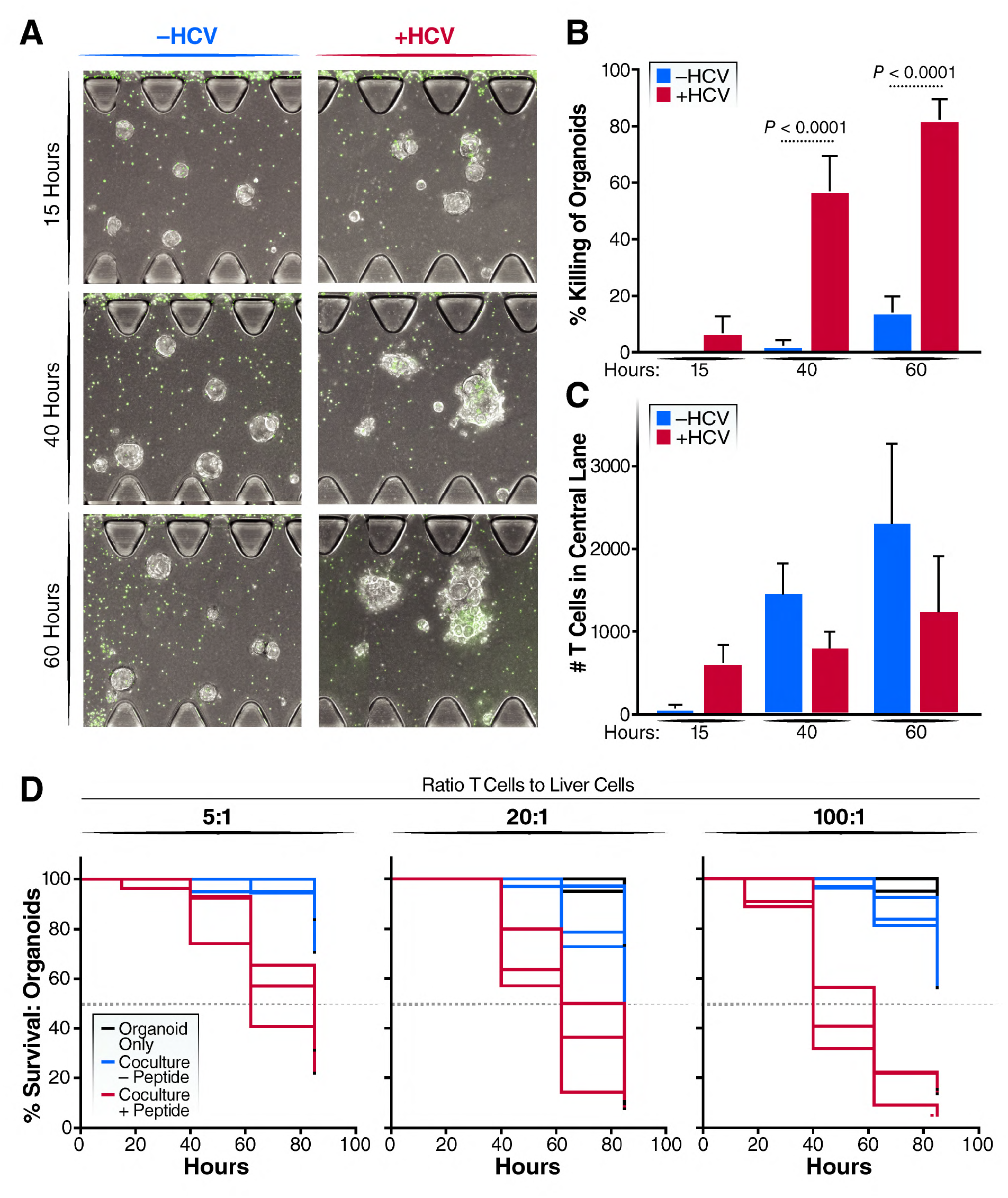
T cell killing response. **(A)** Representative microscopy image of the effect of T cell-liver organoid interaction with and without the HCV peptide presentation on liver organoids. **(B)** Quantification of viability of pulsed and unpulsed organoids at varying time points at a 100:1 effector to target ratio. **(C)** Number of T cells in the central channel in pulsed and unpulsed conditions. (**D)** Survival curve of organoids in monoculture (black), coculture with T cell clones (blue) and coculture with T cell clones after peptide pulsing (red) in 100:1, 20:1 and 5:1 coculture ratio of T cells to liver cells. Each line represents one replicate.

We repeated this experiment with various effector T cell to target organoid ratios. We tested a 100x, 20x and 5x excess of T cells compared to liver organoids in the coculture experiment. Addition of T cells at varying effector to target ratios influenced the level of killing (Fig 5C). Overall, the final percentage of cell death correlated with the number of T cells added into the experiment but a 5:1 effector to target ratio took longer to reach the 50% viability limit. At this ratio, 50% of the peptide-pulsed organoids died only after 80 hours, compared to the 100:1 ratio, which induced 50% cell death at 40 hours (Fig. 5C). A high ratio (100:1) increased the dynamic range between the peptide-pulsed and control co-cultures likely because–unlike in a well-based coculture system– not all T cells access the central channel and not all T cells in the central channel contact the organoids due to the presence of the extracellular matrix, necessitating a higher number of input effector cells.

## Discussion

We report the establishment of a tractable coculture system of liver organoids and antigen-specific T cells, which we envision could be applied towards identifying immunogenic HCV epitopes that lead to T cell-mediated killing of virally infected liver cells. We report here that organoid medium is suitable for coculture of the two cell populations in a microfluidic device that enables the spatial and temporal analysis of individual organoid responses by imaging. Using a novel HCV antigen-specific CD8^+^ T cell clone, we quantify functional T cell activity and specific killing of peptide-presenting liver organoids in the device using common light and fluorescent microcopy equipment and commercially available quantification programs. Parallel work in our lab has shown that ASC-derived liver organoids can be infected with patient-derived isolates of HCV, so we expect this system to provide a new platform to study T:liver cell interactions in a physiological, individualized *in vitro* setting. The system could also prove useful to test effectiveness of chimeric antigen receptor (CAR) T cells in the context of liver infections and liver cancer, and of recently developed neoantigen directing TCR/anti-CD3 fusion proteins^28,29^.

Liver organoids are typically cultured by mixing individual or small aggregates of cells embedded in small droplets of BME. The stochastic, heterogeneous distribution of organoids throughout the relatively large BME droplets makes it challenging to observe the response of individual organoids to infection and presence of T cells. Distribution of organoids in different planes of focus in the droplets make it difficult to compare and contrast individual organoids directly. In addition, the droplet system also lacks ease of registry of individual organoids thereby rendering their longitudinal monitoring inherently challenging. In contrast, studies in the microfluidic chip-based platform enables spatial and temporal observation of individual organoids. The fixed orientation of organoids due to embedment in BME and the monoplanar distribution due to the thin height configuration of the device channels yields a relatively uniform horizontal distribution of organoids. In addition, T cells suspended in culture media can traverse freely through the device and encounter the immobilized organoids. Collectively, this configuration enables longitudinal monitoring and preserves the heterogeneity in biological responses of both the organoids and T cells.

T cells are independently introduced into the system by suspension in culture media, thus mimicking the perfusion of blood entering a tissue. By establishing a lateral pressure gradient across the chip, T cells and media flow in a uni-directional manner through the central channel, which is laden with the matrix-embedded organoids. This spatiotemporal controlled coculture system allows for the selective study of T cells that are able to penetrate the extracellular matrix microenvironment of the central lane and interact with the liver organoids. In addition, parameters such as the ratio of T cells to organoid cells and persistence of interaction between the two cell types can be evaluated. Lastly, the device configuration allows for comparative analysis between T cells that interact with the organoids and the ones that fail to interact and remain in either of the media channels. Collectively, this system can identify specific T cell populations with improved or targeted cellular cytotoxic response towards HCV.

As a model of CD8^+^ T cell response to infection, we chose an immunogenic peptide from the HCV NS3 protein (NS3 coordinates 1406-1415), a key multifunctional enzyme required for viral replication^30^. Analysis of the livers and PBMCs of patients with chronic HCV infection demonstrated that approximately 75% of patients harbored CD8^+^ cytotoxic T cells specific to epitopes located within the NS3 protein^31^. Our study underscores the immunological function that NS3-specific epitopes, here KLVALGINAV, can play in the host response to HCV infection based on the effective cell killing induced by pulsing with the epitopic peptide. Because viral induced liver damage could be mediated both by direct infection of parenchymal cells as well as immune cell cytokine release, the addition of highly reactive immune cells is essential for modeling liver damage during HCV infection. By pairing immune competent cells with adaptive immune cells, we have established a system to explore interventions along the liver cell/immune cell axis. The addition or removal of cytokines, blocking treatments, stimulations of T cells, and other pharmacological interventions could all be readily modeled in this system to attain a better understanding of how such interventions affect liver viability.

Employing a microfluidic chip with distinct compartments for solid tissue and media channels in culture preserves the physiological T cell migration and motility behavior in response to varying chemotactic signals from the liver tissue. Additionally, this compartmentalized culture format allows incorporation of additional relevant immune or liver cell types, such as NK or Kupffer cells, to understand their role in HCV infection. With the re-expanding number of cases of HCV globally and domestically, this novel platform can be used to gain unique insight into HCV immunobiology, potentially accelerating the already long path towards an effective vaccine.

## Materials and Methods

### Liver organoid culture

Liver organoid culture is initiated by 3D culture of EpCAM+ cells freshly isolated from liver resections, using methodology previously described^15^. Briefly, bead-sorted EpCAM+ liver cells were mixed with 50 μl basement membrane extract (BME2) and cultured in a 24-well plate overlaid with basal media (Advanced DMEM with 1% glutamax and 1% pen/strep) containing B-27 (50X, ThermoFisher), N-2, (100X ThermoFisher), 25 ng/ml Hepatocyte Growth Factor (Stemcell Technologies), 50 ng/ml Epidermal Growth Factor (Stemcell Technologies), 10 nM Gastrin I, 1 mM N-Acetylcysteine (Sigma), 100 ng/ml Fibroblast Growth Factor-10 (Stemcell Technologies), 10 μM Forskolin (Stemcell Technologies), 5 μM A83-01 (Tocris Bioscience) and 1:1000 diluted R-spondin-1 conditioned media. Prolonged culture in the above mentioned format generated multicellular liver organoids. Culture media in the reservoir was fully replaced 2x/ week and organoids were split 1:4 with TrypLE digestion every 2-3 weeks.

### Real-time quantitative PCR

RNA was isolated according to manufacturer’s directions with the Qiagen RNeasy mini kit. Organoids were lysed in BME2 with 350 μL of buffer RLT, and RNA was extracted according to the manufacturer’s protocol. The isolated RNA was transcribed into cDNA using oligo(dT)_18_ primers (Thermo Scientific), random hexamer primers (Thermo Scientific), and AMV-reverse transcriptase (Promega). Transcripts were quantified by adding 10 ng of cDNA to a master mix containing 8 pmol of forward and reverse primers, water and 2X SYBR Green Master Mix (Thermo Scientific) to a total of 10 μL. Each assayed gene was run in triplicate for each sample on an Applied Biosciences thermocycler under the following conditions: 50°C for 2 minutes, 95°C for 10 minutes, followed by 40 cycles of 95°C for 5 seconds and 60°C for 30 seconds. Gene expression for the target genes was reported relative to the 18S housekeeping gene.

### Light sheet microscopy

Organoids were fixed for staining according to a previously published protocol^15^. Briefly, organoids were removed from the cell culture plate with ice-cold PBS and then were washed 3x with cold PBS to remove BME2. Organoids were fixed for 30 minutes on ice with 4% paraformaldehyde, washed 3x with cold PBS and stored at 4 °C for up to two months. Before staining, organoids were blocked with PBS supplemented with 0.5% Triton X-100, 1% DMSO, 1% BSA, and 1% donkey serum (blocking buffer) overnight at 4°C. Primary antibody was added at a 1:500 dilution in blocking buffer and incubated at 4°C for 48 hours, and washed 5x with PBS. Secondary antibodies were added in PBS at a 1:250 dilution and incubated at room temperature for several hours. Organoids were washed 5x in PBS to remove secondary antibodies and stained with Hoescht for 10 minutes at 1:1000. Organoids were imaged on Zeiss LSM880 Confocal. Images were processed using a combination of the Zeiss software, ImageJ 1.51f, and Imaris 9.3.

### Peptide pulsing of organoids

Organoids were removed from BME by addition of cold basal media and washed with additional 10 ml of cold basal media. The organoids were resuspended in 200 μL of PBS with 5 μg of either HCV KLVALGINAV peptide, nonspecific peptide, or no peptide, and incubated at 37°C for 1 hour. At the end of the incubation period, organoids were washed 2X in basal media, mixed with BME and seeded into the central lane of the microfluidic chip.

### Generation of CD8^+^T cell clone

CD8^+^ enriched PBMC from HCV spontaneous resolver, SR01, were stained with the A0201-KLVALGINAV-PE tetramer (NIH tetramer Core Facility, Emory University) and CD8-FITC Ab (BD Biosciences, San Jose, CA), and sorted on a FACS Aria cell sorter. Tetramer Sorted CD8^+^ T cell clones were established by limiting dilution seeding of A0201-KLVALGINAV sorted CD8^+^ T cells at either 3, 1 or 0.3 cells/well in 96-well U-bottom plates along with 5 x 10^4^ irradiated allogeneic PBMC feeders, 0.04 μg/ml anti-CD3/ml in RPMI medium supplemented with 10% FBS and 40 u/ml rIL2 (Clone Medium). Plates were cultured at 37°C and 6% CO_2_. Plates were refed every 3-4 days by removing 100 μl of clone medium and replacing with 100 μl fresh clone medium. Between days 7-14, clones displaying growth were transferred into 24-well plates and restimulated with 2 x 10^6^irradiated allogeneic PBMC feeders and 0.4ug anti-CD3/ml in clone medium. Clones were maintained by restimulating every 2-3 weeks with irradiated allogeneic PBMC feeders and anti-CD3.

### CD8^+^T cell culture

CD8^+^ T cell clones were maintained in culture with periodic stimulation (every 2-3 weeks). Briefly, T cell clones were plated in a 24-well plate at 1×10^6^ cells/well in RPMI + 10% FBS + 40u/ml rIL2 (clone medium). Irradiated PBMC feeders (2 x10^6^) from an allogeneic donor were added to each well along with 0.08ug/ml anti-CD3. Plates were incubated at 37°C and refreshed every 3-4 days with fresh clone medium. When wells reached confluency, the clones were pooled together and placed in T75 flasks until the next restimulation. To activate the T cells, they were treated with 0.1mg/ml PMA and 0.5 μM ionomycin while simultaneously treated with BFA and Monensin and incubated for five hours at 37°C.

### Chromium release assay

The 721.221 class I MHC deficient cell line transfected with the A0201 Class I MHC molecule served as targets for the chromium release assay. The A0201 class I transfectant was simultaneously loaded exogenously with peptides at 10 μg/ml concentration and 25 μCi of ^51^Cr for 1 hour at 37°C. The A0201 targets were washed 3X and resuspended at 5 x 10^4^ cells/ml. Recombinant vaccinia virus (rVV) infected A0201 targets were infected at an MOI of 10:1 for one hour, washed 1X and cultured O/N at 37°C. The next day the rVV targets were washed and labeled with 25 μCi of ^51^Cr for 1 hour at 37°C. The clones were seeded at E:T ratios of 20, 5 and 1:1 in 96-well bottom plates. The targets were added to the clones at 5000 cells/well. Chromium release assays were incubated for 3-4 hours at 37°C. 50μl supernatant was harvested from each well and placed in 96-well Luma Plates containing a dry scintillant (Perkin Elmer, Waltham, MA). Dried plates were counted on a MicroBeta counter (Perkin Elmer, Waltham, MA). Percent specific ^51^Cr release was calculated as follows: (Experimental release-Spontaneous release)/ (Maximum release-Spontaneous release) x 100 Responses were considered positive if the percent specific lysis was twice or more above background at 20:1 and 5:1 E:T ratios.

### Fluorescent labeling of T cells

T cells were stained with CellTracker Green CMFDA (ThermoFisher Scientific, C2925) for real-time fluorescence imaging in microfluidic chip. CellTracker staining solution was freshly prepared by diluting the stock (2mM in DMSO) in prewarmed PBS to a final concentration of 10μM. T cells were washed 2X in PBS at 400xg for 5 minutes to remove any traces of cell culture media. Following this, the cells were suspended in the CellTracker staining solution and incubated at 37°C for 30 minutes. Cells were washed 3X to remove unbound dye and resuspended in the appropriate media for the subsequent steps.

### Organoid-T cell coculture in microfluidic chip

Liver organoids and T cells were cocultured in single use, 3D cell culture chips obtained from AIM Biotech (DAX-1). The microfluidic chips consist of a central channel for the 3D culture of cells embedded in hydrogel and two flanking channels for introducing media and/or secondary cell type(s). In this study, liver organoids were cultured in the central channel and the flanking channels were used for cell culture media exchange or for introducing T cells suspended in media. Both sides of the central channel are bordered by vertical posts with a triangular base that prevents leaking of content from the central channel to the flanking channels on either side. Regular gaps between these vertical posts ensure that the interstitial flow of media from the flanking channels towards the central channel can still occur uninterrupted. To initiate the coculture in chips, liver organoids were gently dissociated with TrypLE to obtain organoids of about 75-150 cells each, rinsed 2X in cold basal media to remove any residual BME, and pulsed with HCV or nonspecific peptides as described in the previous section. Organoids were resuspended in the appropriate volume of cold EM to generate a final concentration of approximately 20 organoids/µl. Following this, the organoids suspended in EM were mixed with equal volume of BME2 to bring the final concentration to 10 organoids/µl and placed on ice until loading into chips. 7.5µl of this organoid suspension was injected into the central channel of the 3D chip using micropipettes by carefully pipetting in from either one of the media ports without generating any air bubbles. Chips were inspected under the microscope to ensure the organoid suspension was uniformly dispersed throughout the length of the central channel prior to incubating at 37°C for 15 minutes to allow for complete cross-linking of BME. For the monoculture conditions, the media channels were loaded with EM warmed to 37°C. For the coculture conditions, media channels were loaded with T cells suspended in EM at the relevant cell density. To load only media, 10µl media was carefully pipetted into the left-side media inlet of both top and bottom flanking channels. Following this, the troughs above the left and right side media-inlet ports were carefully filled with 80µl media and 50µl media, respectively. Similarly, the bottom media channels were loaded with 50µl and 40µl media on either side. For the coculture conditions, the required number of CellTracker stained T cells were suspended in 10ul EM and injected into the top flanking media channel only. Cell culture media was replaced every 24 hours by carefully aspirating the media out from the troughs without reaching into the media inlet (Fig. 3A). The microfluidic chips were carefully incubated at 37°C for the entire culture duration. Microfluidic chips along with 3cm petri dishes filled with sterile water were placed in 1-well plates for easy handling and minimization of evaporation of cell culture media from the chips.

### Distinguishing between viable and nonviable liver organoids

Liver organoids were classified as viable or nonviable at different time points of image acquisition based on the morphometric analysis of the organoids. Liver organoids were imaged with phase contrast microscopy through maximum intensity projections of z-stack images obtained across the height of the central channel. The entire area of the central channel was imaged (10.5mm X 1.3mm), and using the spatial coordinates, the different tiles were stitched together using the built-in tiling feature of FIJI image analysis software with an overlap of 5%. At the 0-hour time point, all organoids in the central channel were traced using the elliptical selection tool in FIJI and the morphology parameters were measured and recorded. Aggregates of individual organoids or those attached to the posts or borders of the central channel were excluded from the analysis due to irregular geometries. Organoids that appeared unhealthy at baseline were also excluded from the study. The annotated set of organoids were monitored over time and morphometric changes were analyzed. Organoids with decreasing total area, combined with a sharp decrease in the circularity measure (<0.7) were deemed to be nonviable. Prism software was used for data visualization and statistical analyses.

### Viability assay for CD8^+^T cells

Real-time viability of T cells in different culture conditions was determined using DRAQ7 dye (Abcam, ab109202), which selectively fluorescently stains dead cells. Briefly, cells were cultured in media containing DRAQ7 throughout the duration of the viability assay at a final concentration of 3µM. At each time point of analysis, T cells in 3D chips were imaged using fluorescence microscopy to quantify the total number of DRAQ7+ cells using excitation of 647nm and emission at 681nm. Total live T cell count was obtained through the quantification of CellTracker+ cells and total dead cell count was established through quantification of DRAQ7+ cells.

### Image analysis for T cell viability assay

Fluorescent images with z-stacks were acquired on a Zeiss inverted microscope. For each chip, images were acquired to cover the entire area of the central channel (10.5mm X 1.3mm). All the subsequent image analysis was performed using FIJI image analysis software. Tiles were stitched to recreate the entire field of view of the central channel using the built-in tiling feature of FIJI. Images were split into the different color channels. Total number of live cells was calculated using the CellTracker Green+ cells and total number of DRAQ7+ cells yielded the dead cell count. To count cells, all the images from different conditions were set to the same signal threshold, images were converted to binary format, and the built-in “analyze particles” plugin feature was used to count the numbers of CellTracker and DRAQ7+ cells. At least 3 biological replicates were used for each condition.

---

**Table S1:**
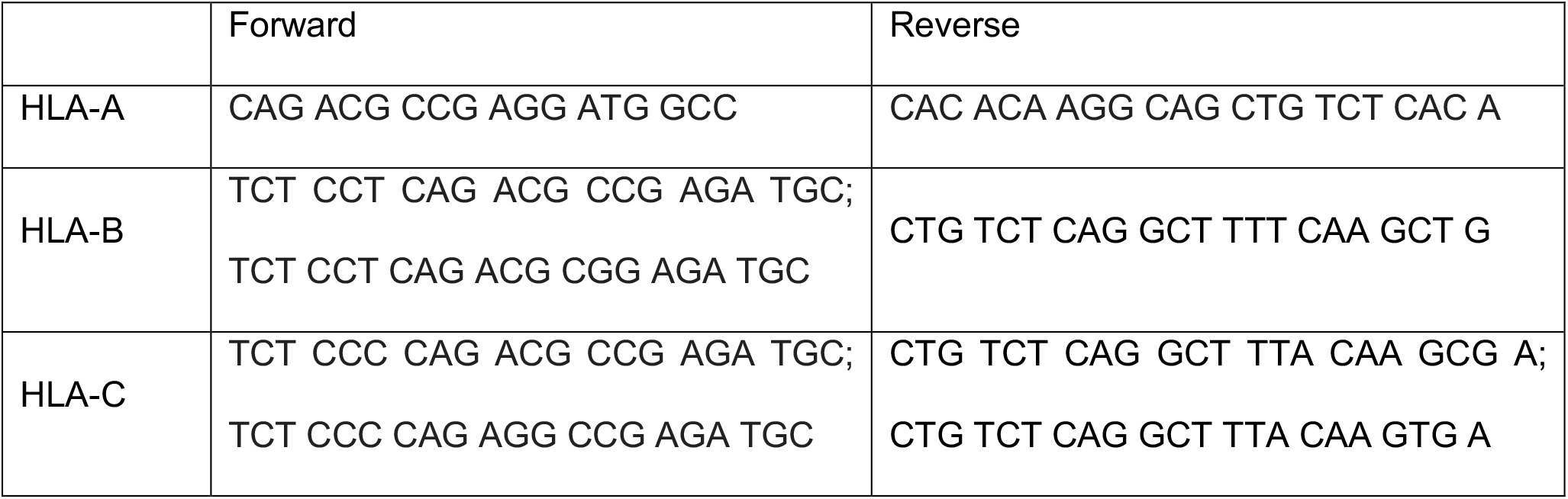
qPCR Primers

**Table S2:** Antibodies

| Antibody | Company | Catalog number | Method |
| --- | --- | --- | --- |
| Anti-B2M | Cell<br>Signals | 12851S | Microscopy |
| Anti-rabbit A594 | Invitrogen | A11012 | Microscopy |
| PerCp-Cy5.5 Anti-IFNγ | BioLegend | 502526 | Flow<br>Cytometry |
| BV421 Anti-CD107a | BioLegend | 328637 | Flow<br>Cytometry |

## Acknowledgments

We thank the Ibrahim El-Hefni Liver Biorepository at the California Pacific Medical Center Research Institute for providing patient tissue samples used to generate liver organoids. We thank John CW Carroll for help with graphic design. We also thank Dr. Kathryn Claiborn for editorial assistance.

